# Search for candidate substances from the spiny red gurnard *Chelidonichthys spinosus*, as novel food additives that color cultivated meat red

**DOI:** 10.1101/2024.07.11.601406

**Authors:** Eriko Shimada, Yusuke Tsuruwaka

## Abstract

The spiny red gurnard *Chelidonichthys spinosus* is a fish with a unique evolution of brightly colored fins consisting of various colors. In Japan, the fish is also used as an ingredient for fish paste products such as fish cake, because of its small edible portion in the processed food industry.

We previously reported on the preparation of “sashimi” in the field of cellular agriculture. In this study, we attempted to explore for cell types suitable for “surimi” or fish paste products and to establish a culture method. In particular, we focused on the development of raw materials for cultivated meat containing pigment cells from edible fish, the spiny red gurnard. As a result, migratory cells were succeeded to be observed from a portion of the fish fin by the migration method. Three types of cells were found in the migrating cells: epithelial-like, fibroblast-like, and those cells containing pigment. Five or more passages of culture were possible, suggesting that the cells could be used as one of the raw materials for cultivated meat surimi in the future. The pigmented cells in particular are expected to be a naturally occurring material that can be used to color surimi.

## Introduction

Lately, increasing attention has been paid to cellular agriculture in terms of a solution to the food shortages in the world because of its ability to provide a sustainable food supply, and has gradually begun to penetrate society (Nature food, 2022; Bomkamp et al., 2023). Cultivated meat can be produced from animal cells in the clean environment of a laboratory, without the need for conventional animal breeding or slaughter. It is anticipated that the accumulation of information on various edible cell types will become very important in the future. Our team is particularly interested in cultivated fish meat. We have successfully implemented thread-sale filefish and scorpionfish cells in society and reported a method for producing cultivated meat ‘sashimi’ from the cells in thread-sale filefish (Tsuruwaka *et al*. 2014; Tsuruwaka &Shimada 2022). In this study, we searched for new candidate raw materials for cultivated meat seafood and focused on the spiny red gurnard, which is distributed on market as a food resource.

The spiny red gurnard *Chelidonichthys spinosus* is a species of sea robin native to the northwestern Pacific Ocean where they occur at depths of from 25 to 615 meters (Carpenter and Niem 1999). This bottom-living fish is characterized by its vermilion body color and bright blue pectoral fins. In Japan which is one of the world’s largest consumers of fish, the gurnards are considered as a tasty fish species and are served as sashimi, grilled, fried, simmered, and in arabesque soup. In the processed food industry, it is also used as an ingredient in fish paste products such as surimi or *kamaboko* in Japanese. We would like to report on the characteristics of the gurnard cells; from the cell culture to its application to bio-cooking.

## Results and Discussion

The outgrowth of the gurnard cells was observed around the fin tissues as soon as those seeded. A rapid migration of the cells from pectoral fins occurred within a few hours of initiation of culture (Fig. 1 and Movie 1). The migratory cells were adherent, small and showed elliptical shape. They seemed to be epithelial-like cells and continued proliferating for a couple of days. The gurnard cells that emerged from the caudal fin tissues showed similar cell morphology to those from the pectoral fin (data not shown).

**Figure 1.**
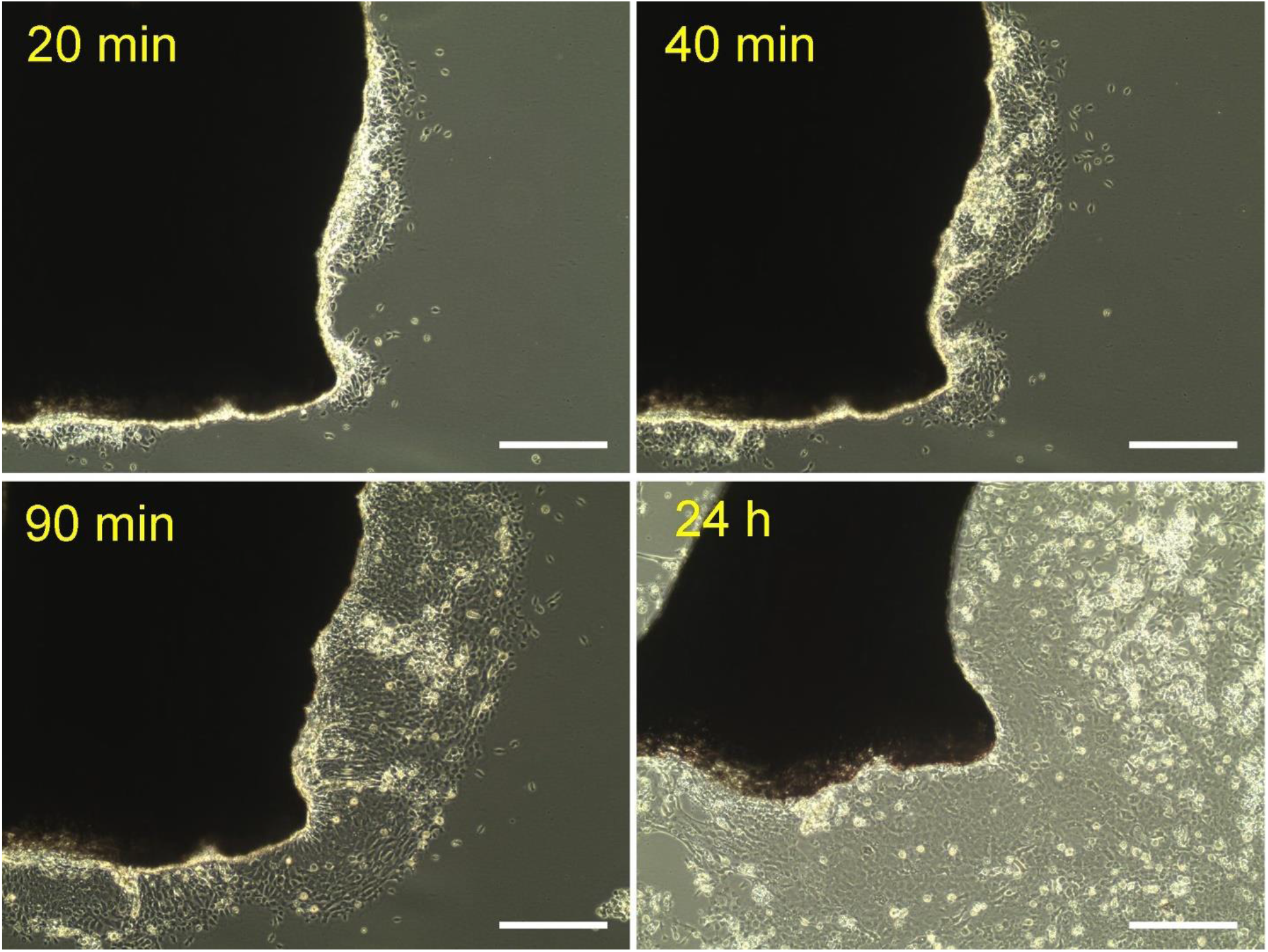
Migratory cells at 20 min, 40 min, 90 min, and 24 h after seeding the fin tissue of *C. spinosus*. Scale bars: 20 µm.

A few days later, fibroblast-like cells then emerged around the tissue and continued to proliferate overwhelmingly the epithelial-like cells (Fig. 2). Eventually, the epithelial-like cells disappeared, which resulted in only the fibroblast-like cells survived in the culture flask (Fig. 2). The fibroblast-like cells were subcultured at five passages. The doubling time was 27.3 h and the cells moved 1 µm/h (Fig. 3).

**Figure 2.**
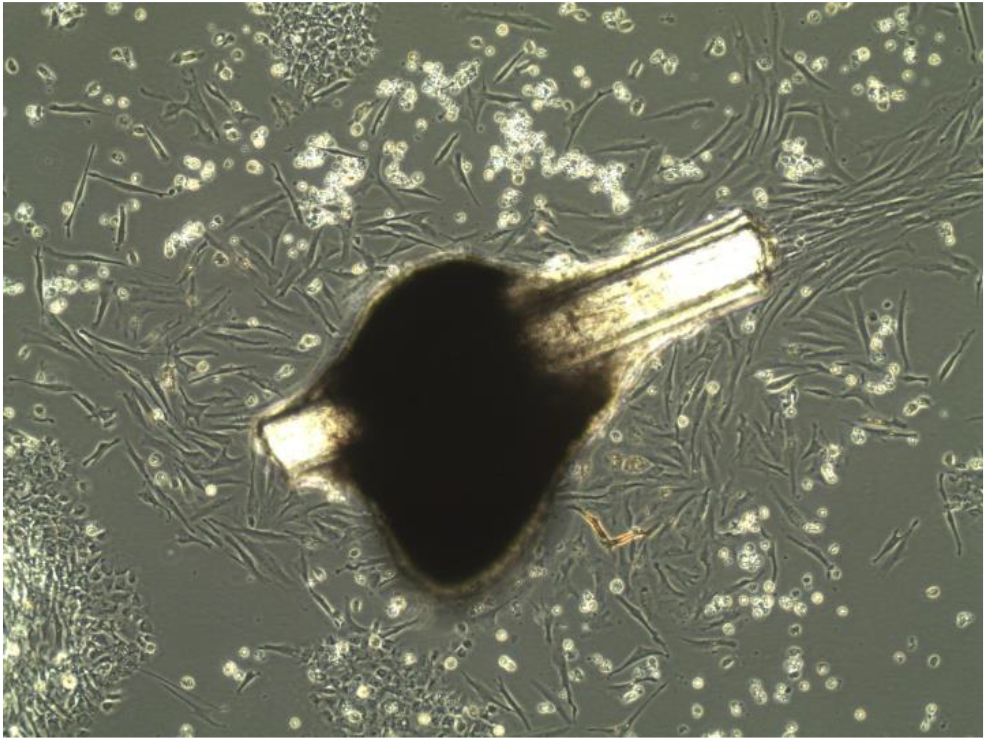
Epithelial-like and fibroblast-like cells of *C. spinosus*.

**Figure 3.**
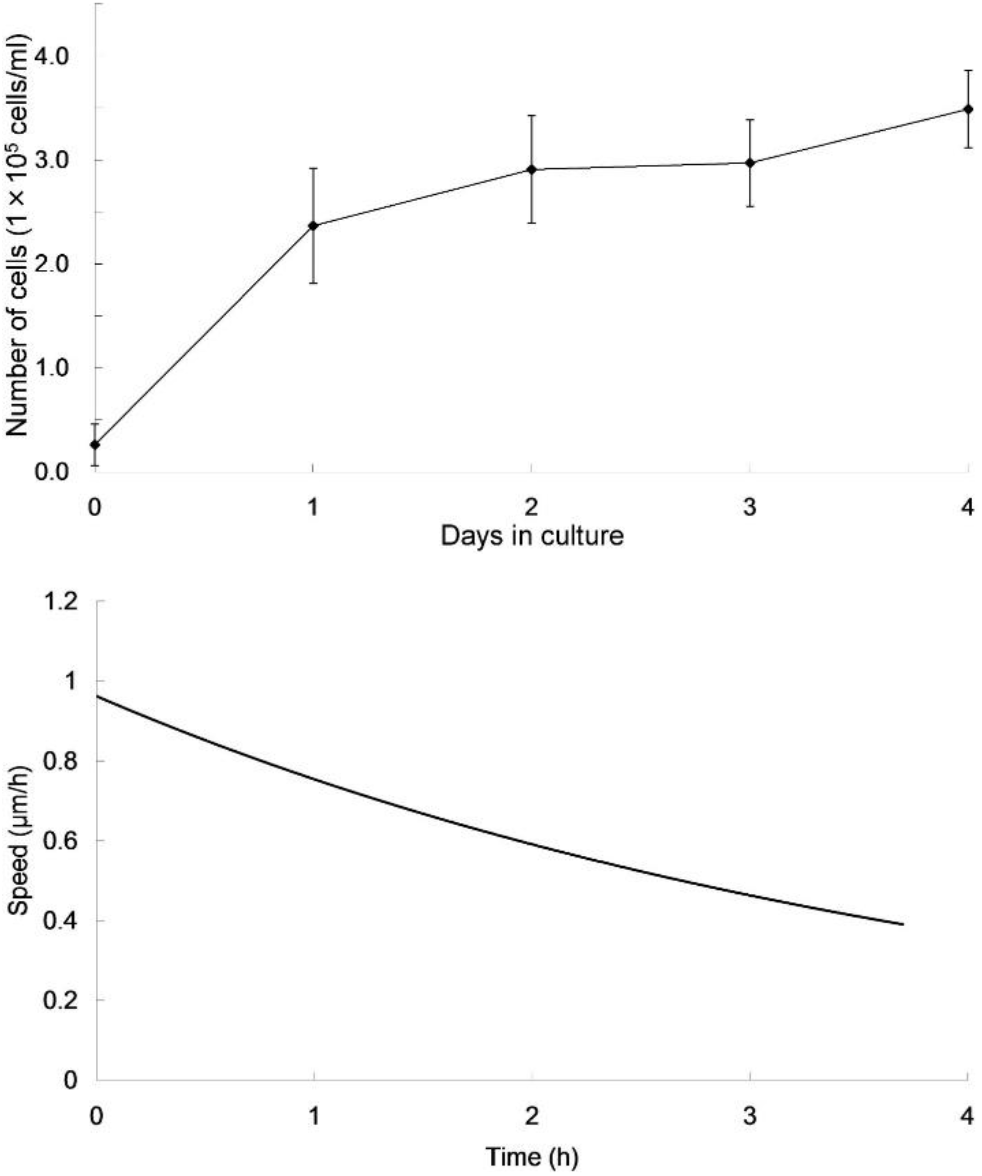
Cell proliferation curve (upper) and velocity curve (lower) of *C. spinosus*.

The gurnard cells were able to be cultured at every temperature we had tested; 12°C, 20°C, 25°C and 30°C. Of those, the cells cultured at 20°C and 25°C looked healthy and proliferating (data not shown).

The gurnard cells were found to be very fast-proliferating and able to migrate at wide range of temperature. These potentials would be beneficial characteristics in cellular agriculture. Of particular interest is the fact that pigmented cells are easily obtained (Fig. 4). That is a property suitable for fish paste products such as *kamaboko*, and could be a tool for using the cells to aid in the coloring of seafood sticks (Fig.5), which originally relied on red food coloring. Providing cultivated meat using gurnards can differentiate its supplier in the marketplace since they produce a novel product line. It can also appeal to consumers looking for sustainable seafood alternatives.

**Figure 4.**
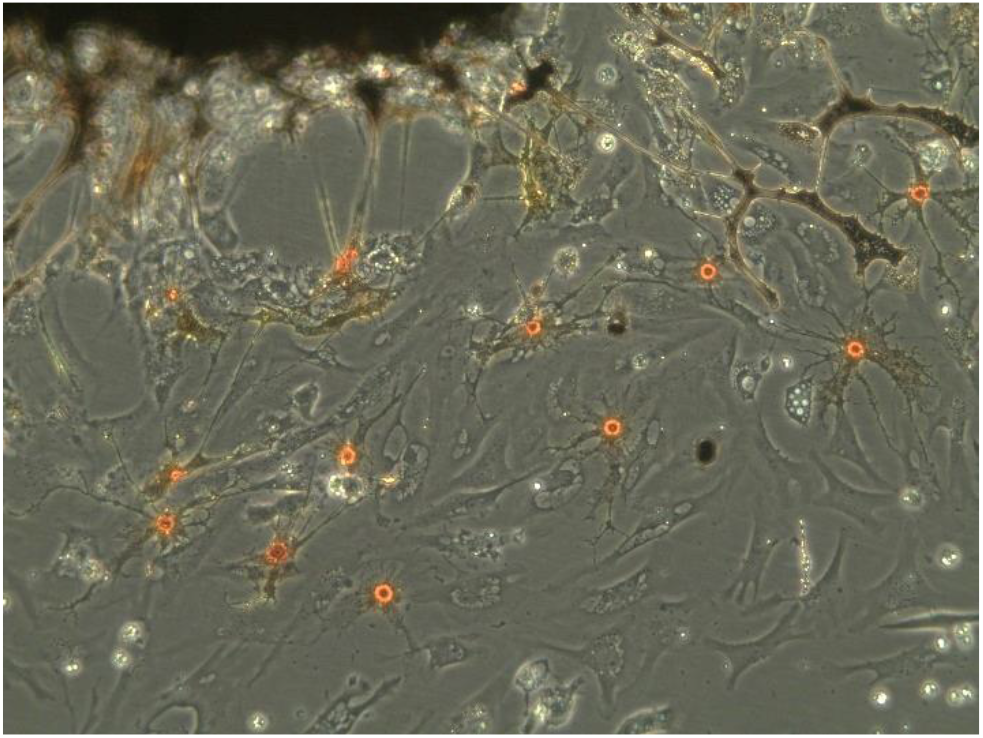
Red pigmented cells of *C. spinosus*.

**Figure 5.**
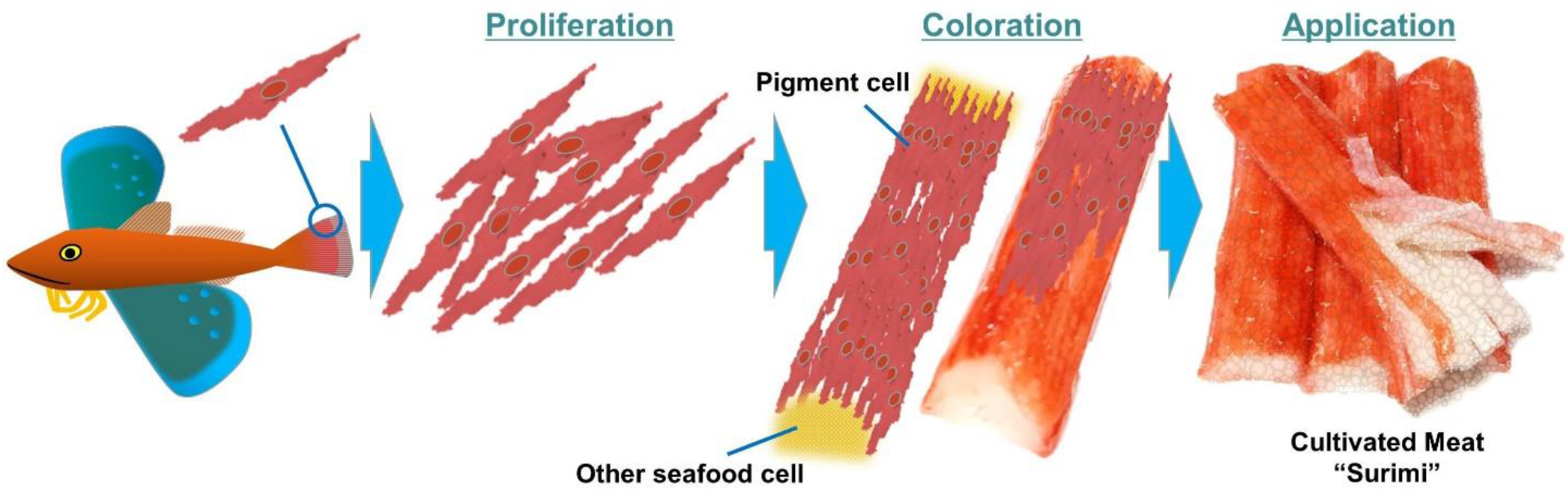
Possible application: color seafood sticks “surimi” with the pigment cells.

## Conclusion

In the present study, we have succeeded in establishing the gurnard’s cell line. Those cells showed the high proliferation rate with the doubling time 27.3 h and the ease of establishment of the cell line. Since the texture of the spiny red gurnard is moderately soft and moist, it would be interesting to mix it with other materials in cellular agriculture as a supplementary tool for processed foods such as “surimi”.

## Materials and methods

### Animal

Live spiny red gurnards *Chelidonichthys spinosus* were captured near Jogashima in Kanagawa Prefecture, Japan, and were maintained in aerated seawater. The salinity and water temperature were maintained at 34.0 ppt and 20±2°C, respectively.

### Development of primary cell culture

Primary cells were prepared the cells as described below. We obtained a 5-mm^2^ cut of pectral and caudal fin tissues from the spiny red gurnard and washed them in water. The tissue was then washed three times with phosphate buffered saline, then six times with penicillin and streptomycin (MP Biomedicals, Santa Ana, CA) on ice. The dorsal fin tissue was cut into 1 mm^2^ in 0.25% Trypsin (MP Biomedicals) 0.02% EDTA (MP Biomedicals) solution and incubated at room temperature for 20 min. The tissue was centrifuged at 1100 rpm and washed twice with Leibovitz’s L-15 culture medium (Life Technologies, Carlsbad, CA). The tissue was immersed in L-15 media containing 10% fetal bovine serum (Biowest, Nuaillé, France) and 1% Zell Shield (Minerva Biolabs, Berlin, Germany), seeded in a 25-cm^2^ collagen I-coated flask (Thermo Scientific, Waltham, MA), and cultured in an incubator at 12-25°C. The culture media was replaced every three days. Cell morphology using an inverted microscope CKX41 (Olympus, Tokyo, Japan) connected to a digital camera ARTCAM-300MI-WOM (Artray, Tokyo, Japan).

The proliferating movie was taken with an inverted microscope Axio Observer. D1 (Carl Zeiss, Oberkochen, Germany) connected to a digital camera AxioCam HR (Carl Zeiss).

The cells were counted using ImageJ software (National Institutes of Health, Bethesda, Washington, D.C. USA). Experiments were repeated five times. Data are presented as the mean ± standard error of the mean (SEM), and differences between groups were identified using Student’s t-test and were considered significant when p < 0.01.

## Acknowledgments

We would like to thank T. Ogawa for technical support.

## Conflict of interest

The authors declare no competing interests.

**Movie 1**. Migratory cells from the fin tissue of *C. spinosus*.

## Notes

### Competing Interest Statement

The authors have declared no competing interest.

